# A multipass membrane protein interacts with the cGMP-dependent protein kinase to regulate critical calcium signals in malaria parasites

**DOI:** 10.1101/2020.07.18.209973

**Authors:** Aurélia C. Balestra, Konstantinos Koussis, Natacha Klages, Steven A. Howell, Helen R. Flynn, Marcus Bantscheff, Carla Pasquarello, Abigail J. Perrin, Lorenzo Brusini, Patrizia Arboit, Olalla Sanz, Laura Peces-Barba Castaño, Chrislaine Withers-Martinez, Alexandre Hainard, Sonja Ghidelli-Disse, Ambrosius P. Snijders, David A. Baker, Michael J. Blackman, Mathieu Brochet

**Author notes:** These authors contributed equally: Aurélia C. Balestra and Konstantinos Koussis.

## Abstract

In malaria parasites, all cGMP-dependent signalling is mediated through a single cGMP-dependent protein kinase (PKG), a major function of which is to control essential calcium signals. However, how PKG transmits these signals in the absence of known second messenger-dependent calcium channels or scaffolding proteins is unknown. Here we identify a polytopic membrane protein, ICM1, with homology to transporters and calcium channels that is tightly-associated with PKG in both *Plasmodium falciparum* asexual blood stages and *P. berghei* gametocytes. Phosphoproteomic analyses in both *Plasmodium* species reveal multiple ICM1 phosphorylation events dependent upon PKG activity. Stage-specific depletion of *P. berghei* ICM1 blocks gametogenesis due to the inability of mutant parasites to mobilise intracellular calcium upon PKG activation, whilst conditional loss of *P. falciparum* ICM1 results in reduced calcium mobilisation, defective egress and lack of invasion. Our findings provide new insights into atypical calcium homeostasis in malaria parasites essential for pathology and disease transmission.

## Introduction

Malaria is an important cause of global morbidity and mortality, accounting for around 405,000 deaths in 2018 (WHO, 2019). The disease is caused by parasites of the genus *Plasmodium*, which possess a remarkable life cycle with multiple cellular differentiation events in both humans and mosquitoes. To sense and respond to changes in these very different environments, malaria parasites use an intracellular communication system that relies on intracellular messengers and protein kinases. Two key signalling molecules in these pathways are cyclic 3’,5’-monophosphate adenosine (cAMP) and 3’,5’-guanosine monophosphate (cGMP), both of which regulate pivotal processes in most eukaryotes (Baker et al., 2017a).

The sole sensor of cGMP in malaria parasites is the cGMP-dependent protein kinase (PKG), which plays vital roles in most stages of the parasite life cycle, including the asexual blood stages that cause all the manifestations of clinical malaria. These roles include control of cellular events required to trigger release of merozoites from red blood cells (RBCs) (Koussis et al., 2020; Taylor et al., 2010) and hepatocytes (Falae et al., 2010), induction of gametogenesis and parasite transmission upon ingestion of gametocytes by a blood-feeding mosquito (McRobert et al., 2008), and sustaining cellular motility necessary for parasite dissemination (Brochet et al., 2014; Govindasamy et al., 2016). A major function of PKG in all these processes is the tightly regulated mobilisation of calcium from intracellular stores within seconds of activation (Brochet et al., 2014; Fang et al., 2018)

In mammalian cells, one of the two PKG isoforms, cGKI (or PKG1), is a well-established regulator of calcium homeostasis. cGKI activation lowers intracellular free calcium upon stimulation by nitric oxide, favouring smooth muscle relaxation or decreasing platelet activation (Francis et al., 2010). cGKI-dependent phosphorylation is known to regulate levels of cytosolic calcium via distinct mechanisms, including either decreased calcium influx by L-type calcium channels (Yang et al., 2007) or sequestration of calcium within intracellular stores. Increased calcium sequestration can be mediated by the phosphorylation of: i) the inositol 1,4,5-trisphosphate (IP_3_) receptor-associated cGMP-kinase substrate (IRAG), which reduces calcium release from intracellular stores by IP_3_-receptors (Schlossmann et al., 2000); or ii) phospholipase-C β3, decreasing IP_3_ generation and subsequent calcium mobilisation (Xia et al., 2001); or iii) phospholamban, which enhances calcium sequestration by the sarcoplasmic reticulum calcium/ATPase pump (Koller et al., 2003). *Plasmodium* parasites are highly divergent from their mammalian hosts (Baldauf, 2003) and no genes encoding clear homologues of IRAG, IP_3_-receptors, phospholamban or L-type calcium channels have been identified in their genomes (Brochet and Billker, 2016).

In the absence from *Plasmodium* of such key signalling molecules, it remains unknown how malaria parasite PKG regulates calcium mobilisation from intracellular stores. We reasoned that the identification of PKG interacting proteins in *Plasmodium* would provide valuable new insights into the role of PKG in calcium homeostasis. Here we identify a multi-pass membrane protein, termed **I**mportant for **C**alcium **M**obilisation-1 (ICM1; Pf3D7_1231400) that interacts tightly with PKG in two different *Plasmodium* developmental stages and species. We show that ICM1 is phosphorylated in a PKG-dependent manner and has an essential function required for intracellular calcium mobilisation downstream of PKG in both the clinically relevant asexual blood stages and in gametocytes that mediate transmission to mosquitoes.

## Results

### *Plasmodium* PKG interacts with ICM1, a multi-pass membrane protein, in both asexual and sexual stages

To identify potential PKG interacting partners in *P. falciparum*, we used a previously established transgenic parasite line expressing PKG fused at its C-terminus to a triple HA tag (Hopp et al., 2012) (PfPKG-HA). PfPKG-HA was immunoprecipitated from extracts of mature schizonts using anti-HA-conjugated magnetic beads and the eluates analysed by quantitative tandem mass spectrometry. The most enriched protein co-immunoprecipitated with PfPKG-HA was identified with high confidence as Pf3D7_1231400, annotated as a putative amino acid transporter (www.PlasmoDB.org), but henceforth referred to as **I**mportant for **C**alcium **M**obilisation 1 or ICM1 (Fig. 1a and Supplementary Table 1). To corroborate these findings, we generated a second transgenic *P. falciparum* line expressing a GFP-tagged PKG (PKG-GFP) (Supplementary Figure 1a-e). Mass spectrometric analysis of pull-downs from extracts of these parasites using anti-GFP nanobodies similarly identified PfICM1 as a top hit (Supplementary Figure 1f and Supplementary Table 1).

**Figure 1.**
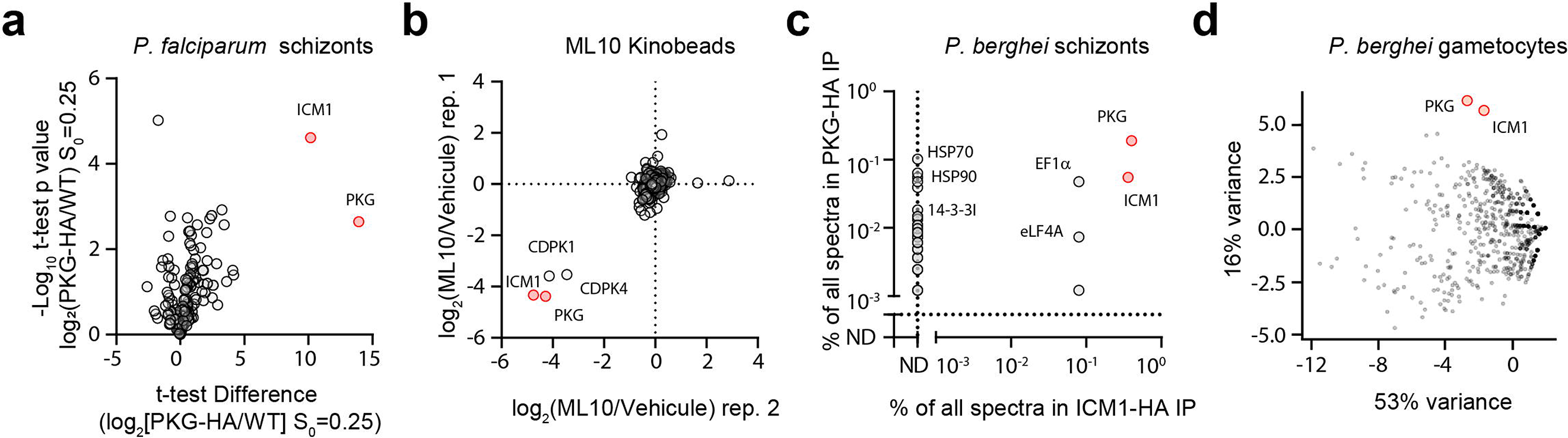
PKG interacts with ICM1 in *P. falciparum* and *P. berghei* schizonts and in *P. berghei* gametocytes. **a)** Volcano plot depicting relative abundance of proteins identified by quantitative mass spectrometry in pull-downs from: *P. falciparum* PKG-HA compared to WT (control) parasites (n=3). Significance (Student’s t test) is expressed as log_10_ of the p value (y-axis). X-axis, enrichment of interaction partners in the *P. falciparum pkg-ha* pull-down compared to controls. **b)** Competitive chemoproteomic studies using Kinobeads on extracts of *P. falciparum* schizonts. **c)** Mass spectrometric quantitation of proteins immunoprecipitated from extracts of *P. berghei* PKG-HA3 (n=1) and ICM1-HA3 (n=1) schizonts. ND = not detected **d)** Mass spectrometric spectral count values for proteins co-purifying with PbPKG-HA3 (n=2) and PbICM1-HA3 (n=2) gametocytes following immunoprecipitation, and displayed in first and second principal components compared with previously published immunoprecipitations of CRK5, CDKrs, SOC2, CDPK4, SOC1, SOC3 and MCM5. See also Figure S1.

To independently validate the putative interaction between ICM1 and PKG, we took an orthogonal chemical proteomics approach. A potent PKG inhibitor (Baker et al., 2017b) (ML10) was used on a Kinobead matrix (Bantscheff et al., 2007) to identify the repertoire of *P. falciparum* proteins targeted by this compound. Proteomic profiling showed that ML10 prevented the binding of just four parasite proteins to kinase inhibitors; these were PfPKG, two calcium-dependent protein kinases (CDPK1 and CDPK4), and PfICM1. The selectivity of ML10 for *Plasmodium* PKG exploits the presence of a small threonine gatekeeper residue in the enzyme ATP binding pocket (Baker et al., 2017b). As both CDPK1 and CDPK4 also possess small serine gatekeeper residues (Tewari et al., 2010), it is likely that ML10 binds these two kinases directly. However, identification of a non-kinase hit in the Kinobead experiments likely reflects either off-target effects of the compound or capture of kinase-associated proteins. These results, combined with the pull down data fully support a direct interaction between PKG and ICM1 (Fig. 1b and Supplementary Table 1).

To interrogate whether the interaction between PKG and ICM1 is conserved across *Plasmodium* species and developmental stages, we exploited a previously-described transgenic *P. berghei* line expressing HA3-tagged PKG (PbPKG-HA3) (Brochet et al., 2014) to immunoprecipitate PbPKG-HA3 from extracts of schizonts and gametocytes. Quantitative mass spectrometry showed that in both stages, PbICM1 (PBANKA_1446100) was the most enriched protein co-immunoprecipitated with PbPKG-HA3 (Fig. 1c and d; Supplementary Table 1). To further validate these results in a reciprocal manner, we generated a PbICM1-HA3 line (Supplementary Figure 1g). No signal was observed for PbICM1-HA3 in western blot analysis of schizont or gametocyte extracts from this line, possibly due to low abundance of PbICM1 or inefficient transfer of the protein (predicted MW ~210 kDa) to membranes. However, a weak HA-specific immunofluorescence (IFA) signal was consistently observed which showed partial co-localisation with the endoplasmic reticulum marker BiP (Supplementary Figure 1h-i). Nevertheless, immunoprecipitation of PbICM1-HA3 from extracts of both schizonts and gametocytes recovered multiple peptides, identified by mass spectrometry to encompass up to 28% coverage of PbICM1. As expected based on the PbPKG immunoprecipitations, PbPKG was the most enriched protein co-immunoprecipitated with PbICM1-HA3 in both stages (Fig. 1c and d; Supplementary Table 1). Of note, no peptides from PbCDPK1 or PbCDPK4 were recovered from either immunoprecipitate, confirming that these kinases do not interact with PbPKG or PbICM1 as previously suggested (Fang et al., 2018)

To further characterise the specificity of PbPKG and PbICM1 co-immunoprecipitations in gametocytes, we compared the relative abundance of proteins recovered in both immunoprecipitates with those of seven other proteins involved in gametogenesis (Balestra et al., 2020). Nine-dimension principal-component analysis (PCA) of all detected proteins confirmed the clustering of PbPKG and PbICM1, consistent with the formation of a complex (Fig. 1d). Taken together, these results demonstrate a conserved, high affinity and exclusive interaction between ICM1 and PKG in at least two developmental stages of two different *Plasmodium* species.

### Modelling of ICM1 shows resemblance to transporters and channels

ICM1 is a multi-pass membrane protein annotated as a putative amino acid transporter. A single ICM1 orthologue is present in all sequenced *Plasmodium* genomes, but no orthologues are evident in non-apicomplexan genomes. The N-terminal segment, which encompasses two conserved clusters of 5 and 4 transmembrane domains (TMDs) respectively, shows high levels of identity between the *P. falciparum* and *P. berghei* orthologues (Supplementary Figure 2). This region is separated from a predicted C-terminal TMD (TMD10) by ~1100 amino acid residues that are less conserved between the two species. Homology modelling using Phyre2 (Kelley et al., 2015) suggests weak similarity of the ICM1 N-terminus to cation transporters, amino acid transporters or IP_3_ receptors (Fig. 2a). To investigate this further, the Iterative Threading ASSEmbly Refinement (I-TASSER) approach was used to model the N-terminal 1500 residues of *P. falciparum* ICM1. Following structure assembly simulation, the Protein Data Bank (PDB) was screened for proteins with the closest structural similarity to the model. Remarkably, the top PDB hit was an IP_3_ receptor (rat cerebellum InsP_3_R1, PDB:6MU1) with a TM (template modelling)-score of 0.967, a RMSD (Root Mean Square Deviation) of 2.06 Å and a coverage of 98.7%, although amino acid identity between the two proteins was low (8.4%) (Fig. 2b). Collectively, these findings suggest that ICM1 may represent a structurally divergent polytopic membrane protein with architectural similarities to mammalian channels, transporters and IP_3_ receptors.

**Figure 2:**
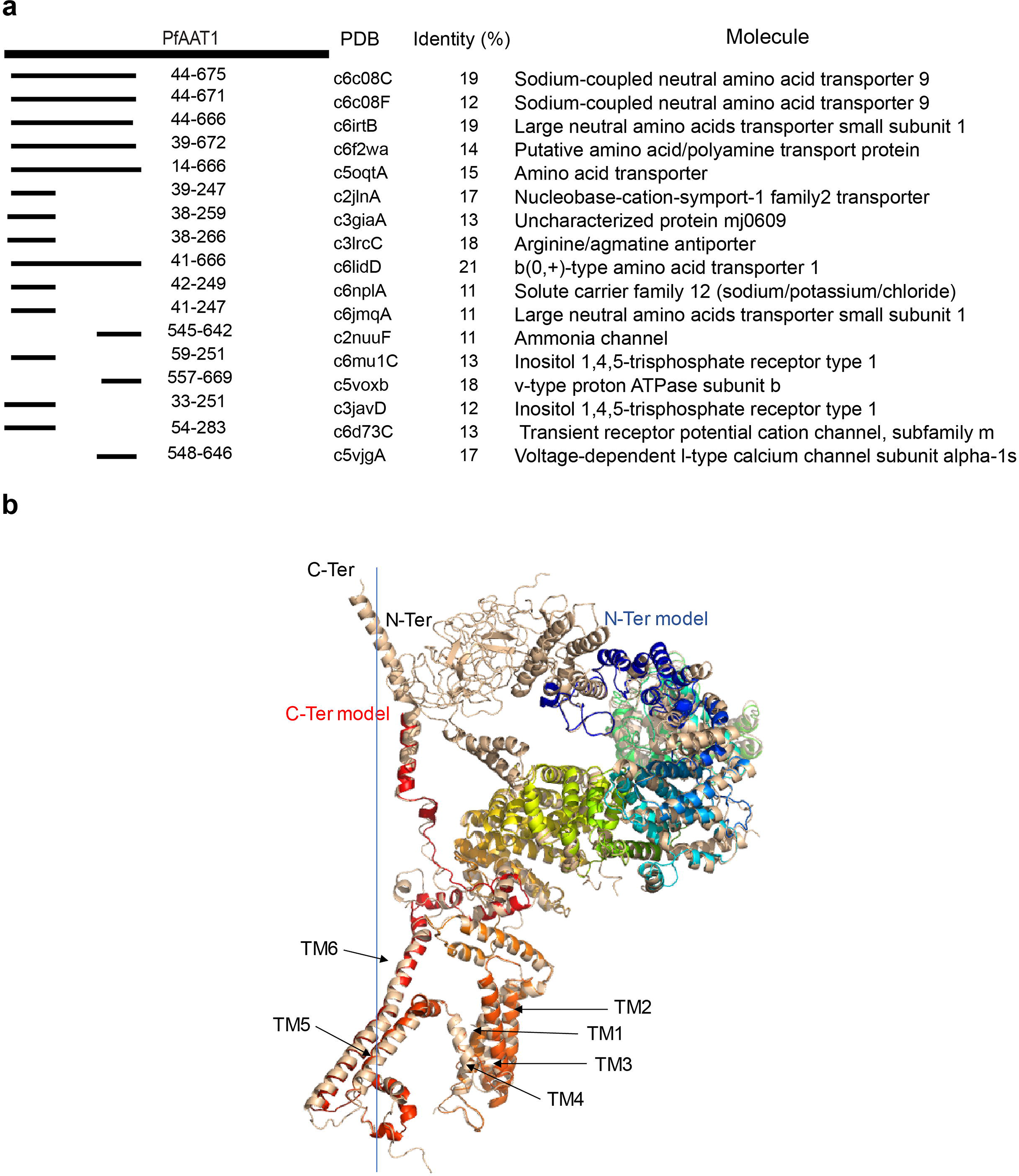
Modelling of PfICM1. **a)** Top hits from PfICM1 homology modelling by Phyre2 indicating the region of homology and the associated percentage of identity, and the molecule description. **b**) Cartoon representation of the I-TASSER AAT1 model (rainbow color) superimposed to its closest structural PDB template (6MU1: InsP3R1 from rat cerebellum, cryo-EM, 4.1 Å, wheat color). The best model was selected with a C-score=−0.30 (C-score range [−5,2], the higher the score, the higher confidence in the model), an estimated TM-score=0.67±0.12 and an estimated RMSD=10.6±4.6Å. The N and C-terminal extremities are indicated and the TM regions are labelled based on the 6MU1 helices topology. The four-fold axis is represented by the vertical dashed line. Only one monomer of the 6MU1 tetramer is represented for clarity. See also Figure S2.

### ICM1 is phosphorylated in a PKG-dependent manner

The physical interaction between PKG and ICM1 raised the possibility of PKG regulating the function of ICM1 through phosphorylation. Several global phosphoproteome studies in *Plasmodium* have identified multiple phosphorylation sites between TMD9 and TMD10 in both *P. falciparum* ICM1 (Alam et al., 2015; Solyakov et al., 2011; Treeck et al., 2011) and PbICM1 (Brochet et al., 2014; Invergo et al., 2017). In a previous study, pharmacological inhibition of PKG activity in mature schizonts did not detect any differential ICM1 phosphorylation under the tested conditions (Alam et al., 2015). However, a phosphoproteome analysis of *P. berghei* guanylyl cyclase beta (GCβ) mutant ookinetes, in which PKG is down-regulated (Moon et al., 2009), showed that PbICM1-S1288 was significantly less phosphorylated in this transgenic line (Brochet et al., 2014). To shed more light on PKG-dependent phosphorylation of ICM1, we profiled the PKG-dependent phosphoproteomes in both gametocytes and schizonts. For the former, we examined phosphorylation events just 15 seconds after activation of gametocytes in the presence or absence of a selective PKG inhibitor (Fang et al., 2018; Green et al., 2015) (Compound A), as PbPKG activity during gametogenesis is only required during the first 15 seconds following stimulation by xanthurenic acid (XA) (Brochet et al., 2014; Fang et al., 2018). In the case of *P. falciparum* schizonts, we generated a conditional knockout of PKG to investigate the phosphoproteome changes in PKG-null mutants. Treatment of this line, termed *pfpkg:cKO*, with rapamycin (RAP) results in disruption of the *pfpkg* gene, expression of mCherry and complete arrest of parasite replication due to the requirement of PKG for merozoite egress (Koussis et al., 2020) (Supplementary Figure 3 and Supplementary movie 1).

Given the low abundance of PbICM1 in gametocytes, we used SDS-PAGE fractionation to enrich for proteins of molecular mass between 180-220 kDa. Analysis of these proteins identified 957 phosphopeptides mapping onto 610 proteins (Supplementary Table 2). Upon pharmacological inhibition of PbPKG, 410 phosphopeptides were less phosphorylated, a large response amplitude that probably reflects the early requirement for PbPKG during gametogenesis (Fig. 3a). Conditional disruption of PfPKG had a similarly profound effect on the *P. falciparum* global schizont phosphoproteome, with more than 1,000 phosphopeptides being significantly hypophosphorylated (Fig. 3b and Supplementary Table 3). GO term enrichment analysis of the proteins showing decreased phosphorylation in *P. berghei* gametocytes suggested deregulation of both S- and M- cell cycle phases, thought to be initiated by calcium mobilisation (Fig. 3c). It is likely that pathways related to cell cycle progression are indirectly controlled by multiple phosphorylation cascades downstream of PKG, including CDPK4-dependent regulation of cell cycle entry (Billker et al., 2004; Fang et al., 2017; Invergo et al., 2017). A similar analysis of the hypophosphorylated proteins in *P. falciparum* schizonts highlighted potential defects in RNA processing and splicing (Fig. 3d and Supplementary Table 3). Interestingly, 11 and 23 proteins predicted to be involved in ion transport were less phosphorylated upon PbPKG and PfPKG inhibition respectively, suggesting tight regulation of ion homeostasis upon PKG-dependent activation. We previously found cGMP-dependent phosphorylation of enzymes involved in the metabolism of inositol phospholipids in *P. berghei* ookinetes (Brochet et al., 2014; Fang et al., 2018). We here confirm these observations for both gametocytes and schizonts, further strengthening the evidence for close interplay between PKG, calcium and phosphoinositide metabolism.

**Figure 3.**
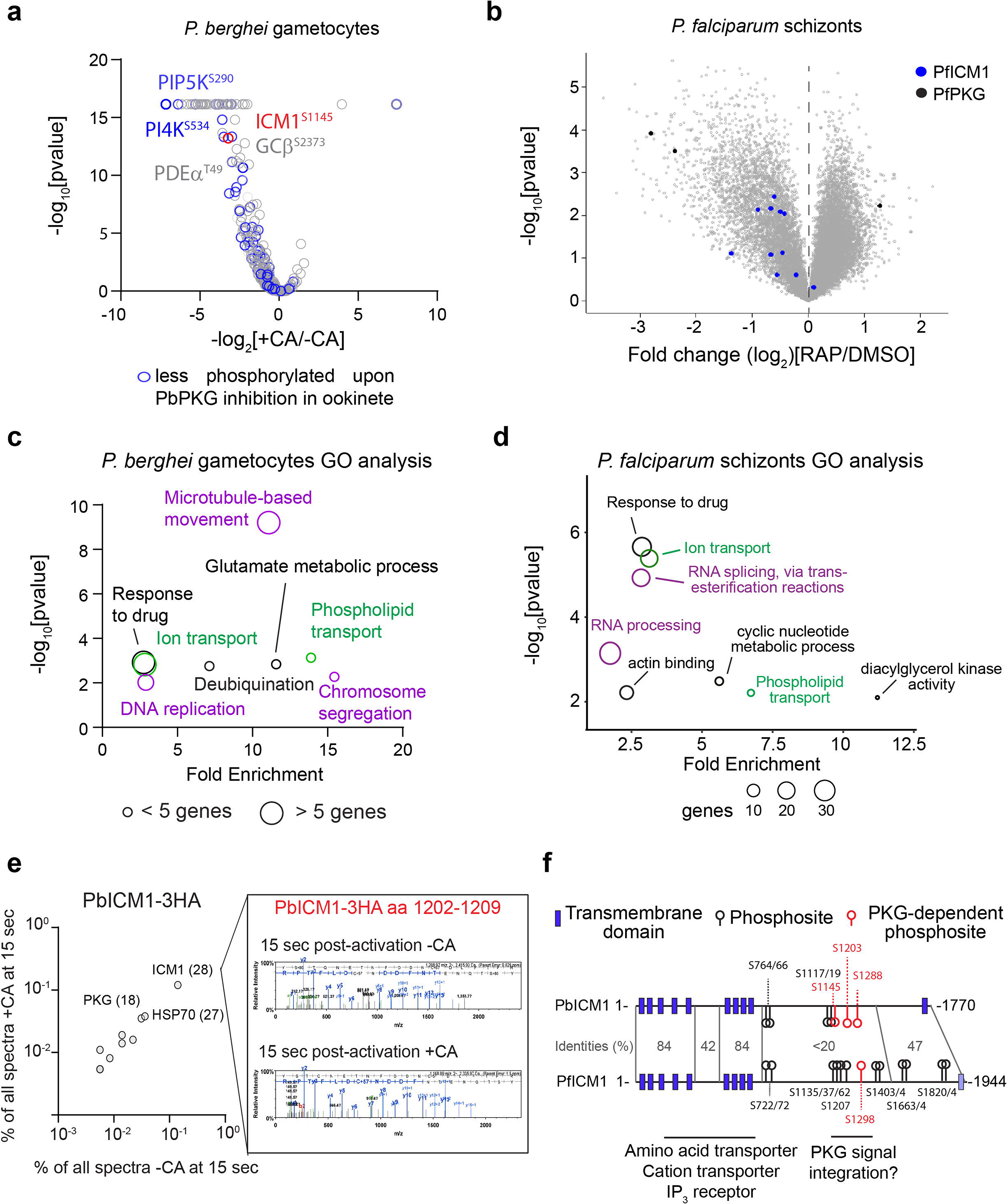
ICM1 is phosphorylated in a PKG-dependent manner. **a)** Volcano plot showing the extent of differentially phosphorylated peptides in *P. berghei* gametocytes 15 sec after XA stimulation in presence or absence of Compound A (CA) (n=3). **b)** Volcano plots showing differentially phosphorylated peptides in DMSO and RAP-treated *pfpkg:cKO* schizonts (n=5). Blue dots correspond to identified PfICM1 phosphosites and black dots to PfPKG phosphosites. **c-d)** GO term enrichment analysis for down-regulated phosphopeptides in 15 sec activated gametocytes upon inhibition of PbPKG by Compound A and of PfPKG-null schizonts. For the latter a cut-off of welch Difference <−1 and >1 with a p value <0.05 and a localisation probability of >0.7 represents a significantly regulated peptide. Representative terms are shown (for a full list see of GO term enrichment analysis in PfPKG-null schizonts see Sup. Table 3) **e)** Relative abundance of proteins co-precipitated with PbICM1-HA3 in 15 sec activated gametocytes in presence or absence of Compound A. Inset, representative spectra indicating that phosphorylation of PbICM1-S1203 is only detected in absence of Compound A. **f)** Schematic of PbICM1 and PfICM1 indicating predicted TMDs and phosphosites detected in this and previous studies. See also Figure S3.

We then focused our attention on ICM1 phosphorylation. Twelve PfICM1 phosphorylation sites were mapped in *P. falciparum* schizonts (Fig. 3b). Ten sites were hypophosphorylated in the absence of PfPKG, suggesting a general defect in phosphorylation of PfICM1 although only two sites (S1289 and S1875) exhibited a fold change of more than 1.8. In gametocytes, intensities of the eight non-phosphorylated peptides mapped onto PbICM1 were not affected by Compound A, indicating that PbICM1 abundance is not dependent on PbPKG activity. We identified two PbICM1 phosphosites, of which S1145 was significantly down-regulated upon PbPKG inhibition. Phosphorylation of PbICM1-S1145 was previously shown to be up-regulated within seconds of gametocyte activation by XA (Invergo et al., 2017) and so possibly represents an important event in PbICM1 regulation. We additionally interrogated the phosphorylation status of immunoprecipitated PbICM1-HA3 in gametocytes 15 seconds after activation, in the presence or absence of Compound A (Fig. 3e). Treatment with compound A had no effect on the interaction between PbPKG and PbICM1-HA3 or on the obtained coverage of PbICM1-HA3. However, S1203 was the only residue detected as phosphorylated following gametocyte activation by XA and no phosphorylation of PbICM1-HA3 was detected upon inhibition of PbPKG. This suggests that ICM1-S1203 is also phosphorylated early during gametogenesis in a PKG-dependent manner.

Collectively, our results indicate that the phosphorylation of at least five ICM1 residues depends on PKG activation. Interestingly, four of the detected PKG-dependent phosphorylation sites are clustered in a region encompassing residues Pf1162 (Pb1145) to Pf1357 (Pb1288) suggesting that this represents an important region for regulation of ICM1 function (Fig. 3f).

### Stage-specific knockdown of PbICM1 reveals a crucial role in early calcium mobilisation required to enter gametogenesis

To study the function and essentiality of PbICM1 during gametogenesis, we first attempted to disrupt the *pbicm1* gene. The gene was previously shown to be resistant to disruption in a global gene knockout study (Bushell et al., 2017), a finding we confirmed here with four unsuccessful disruption attempts using highly efficient PlasmoGEM vectors (Pfander et al., 2013; Pfander et al., 2011). We then tagged endogenous ICM1 with an auxin-inducible degron (AID) coupled to an HA epitope tag (Supplementary Figure 4a) to allow degradation of the fusion protein in the presence of auxin in a strain expressing the Tir1 protein (Philip and Waters, 2015). Addition of auxin to PbICM1-AID/HA gametocytes did not lead to any defect in male gametogenesis. However, we could detect no western blot signal for PbICM1-AID/HA, so were unable to confirm successful degradation of the protein upon auxin treatment (Supplementary Figure 4b). We thus opted for a stage-specific knockdown of PbICM1 by placing the endogenous *pbicm1* gene under the control of the *pbama1* promoter, which is active in schizonts but virtually silent in gametocytes (Sebastian et al., 2012). A P_*ama1*_ICM1 clone was readily obtained and RT-qPCR analysis confirmed a two-fold reduction in *pbicm1* expression in purified P_*ama1*_ICM1 gametocytes (Fig. 4a).

**Figure 4:**
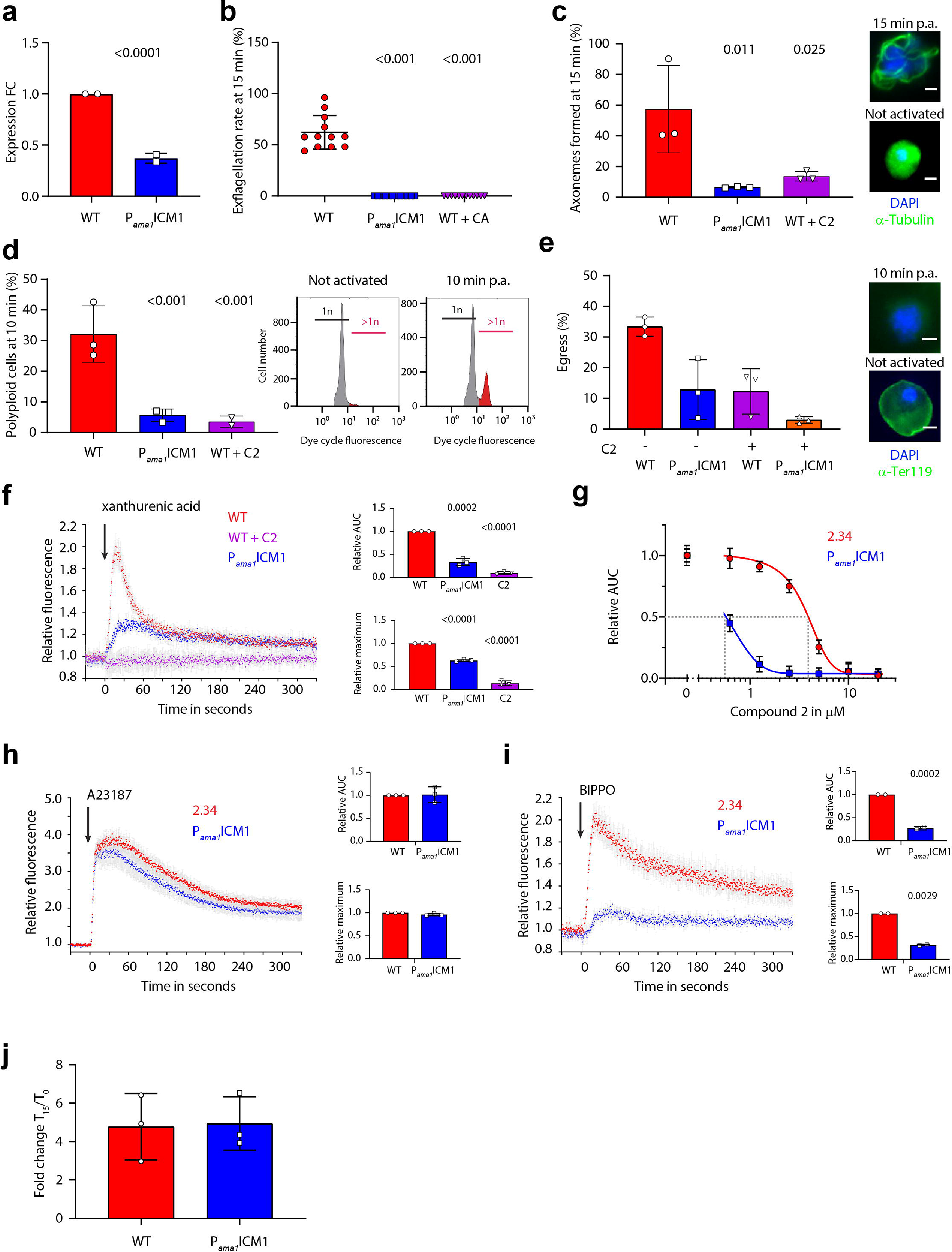
Stage-specific knockdown of PbICM1 partially phenocopies PKG inhibition. **a)** Relative levels of *pbicm1* mRNA in WT and gametocytes (error bars, ± S.D.; 2 independent biological replicates, two-tailed unpaired t-test). **b)** Stage-specific knock-down of PbICM1 leads to a profound defect in exflagellation (error bars, ± S.D.; technical replicates from 3 independent infections; two-tailed unpaired t-test). **c)** P_*ama1*_ICM1 male gametocytes show a strong reduction in fully assembled axonemes 15 min following activation (error bars, ± S.D.; 2 independent infections; two tailed unpaired t-test). Inset pictures show representative tubulin distribution patterns (green), observed in male gametocytes counter stained with DAPI (blue); scale bar, 2 μm. **d)** Reduction in the number of P_*ama1*_ICM1 male gametocytes replicating their DNA. Proportion of male gametocytes undergoing DNA replication was determined at 1 min post-activation (p.a) and is expressed as a percentage of polyploid (>1N) cells (error bars, ± S.D.; 3 independent infections; two tailed unpaired t-test). Insets show the gating used for the flow cytometry analysis. **e)** Egress from RBCs was quantified by flow cytometry based on the presence of the RBC membrane marker Ter-119 in gametocytes, 15 min post-activation. Insets, IFA of WT and P_*ama1*_ICM1 gametocytes activated for 15 min. Green: Ter119; blue: DAPI. Scale bar, 2 μm. **f)** Fluorescence response kinetics of gametocytes loaded with Fluo4-AM upon stimulation with 100 μM XA at t◻=◻0 sec. Insets show quantifications of the relative maximum intensity or area under the curve (error bars, ± S.D.; 3 independent replicates; two tailed unpaired t-test). **g)** Relative calcium response to 100 μM XA, as determined by the area under the curve, of WT and P_*ama1*_ICM1 gametocytes upon treatment with various concentrations of compound 2. **h)** Fluorescence response kinetics of gametocytes loaded with Fluo4-AM upon stimulation with A23187 at t◻=◻0 sec. Insets, quantification of the relative maximum intensity or area under the curve (error bars, ± S.D.; 3 independent replicates; two tailed unpaired t-test). **i)** Fluorescence response kinetics of gametocytes loaded with Fluo4-AM upon stimulation with BIPPO at t◻=◻0 sec. Insets, quantification of the relative maximum intensity or area under the curve (error bars, ± S.D.; 2 independent replicates; two tailed unpaired t-test). **j)** cGMP levels in WT and P_*ama1*_ICM1 gametocytes upon stimulation with BIPPO. See also Figure S4.

To determine the requirement for PbICM1 during gametogenesis, we then compared the effect of *pbicm1* down-regulation with PKG inhibition. Although P_*ama1*_ICM1 gametocytes were microscopically indistinguishable from their WT counterparts, they did not form active exflagellation centres upon treatment with XA, phenocopying PKG-specific inhibition by Compound A (Fig. 4b). A similar result was observed when *pbicm1* was placed instead under the control of the *clag9* promoter, which similarly downregulates *pbicm1* in gametocytes (Supplementary Figure 4c to e). Additionally, PbICM1 down-regulation or PbPKG inhibition both led to significant decreases in axoneme formation, egress from the host RBC, and DNA replication, indicating an early requirement for PbICM1 in male gametogenesis (Fig. 4c to 4e).

We previously showed that PKG is critical for mobilisation of calcium from intracellular stores within seconds of gametocyte stimulation by XA (Brochet et al., 2014). Consistent with a functional link between PbPKG and PbICM1, a strongly attenuated calcium mobilisation was consistently observed in P_*ama1*_ICM1 and P_*clag9*_ICM1 gametocytes (Fig. 4f and Supplementary Figure 4f). The residual signal observed in P_*ama1*_ICM1 gametocytes could possibly be due to incomplete silencing of *icm1* or due to PbICM1-independent pathways downstream of PbPKG also involved in calcium mobilisation. Interestingly, the residual calcium mobilisation observed in P_*ama1*_ICM1 gametocytes was 10 times more sensitive to PKG inhibition by compound 2 compared with WT parasites, further strengthening the evidence for a functional interplay between PbPKG and PbICM1 (Fig. 4g). Importantly, P_*ama1*_ICM1 and P_*clag9*_ICM1 gametocytes responded normally to the calcium ionophore A23187, indicating similar levels of mobilisable intracellular calcium in P_*ama1*_ICM1 and WT gametocytes (Fig. 4h and Supplementary Figure 4g).

To define the functional requirement for PbICM1 in the first seconds of gametogenesis, we took advantage of a phosphodiesterase (PDE) inhibitor, BIPPO, which elevates levels of cGMP in *Plasmodium* by preventing its degradation (Howard et al., 2015). As expected, elevation of cGMP by BIPPO led to calcium release in WT gametocytes. In stark contrast, the calcium response to PDE inhibition was strongly reduced in the P_*ama1*_ICM1 line (Fig. 4i and Supplementary Figure 4h). However, chemical inhibition of PDEs and XA stimulation induced comparable cGMP levels in both WT and P_*ama1*_ICM1 gametocytes indicating that PbICM1 is not required to control cGMP homeostasis upstream of PKG (Fig. 4j). Together, these results indicate that PbICM1 is required for PKG-dependent mobilisation of intracellular calcium.

### Loss of PfICM1 in asexual blood stages inhibits calcium mobilisation leading to inefficient egress and invasion

To study the function of PfICM1 and its link to PKG function in *P. falciparum* asexual blood stages, we investigated the phenotypic consequences of PfICM1 disruption. Since *pficm1* was predicted to be essential (Zhang et al., 2018) we decided to disrupt the gene conditionally using the RAP-inducible DiCre system (Perrin et al., 2018). Generation of the *pficm1:cKO* line was achieved in two steps using a marker-free Cas9-mediated strategy. In the first of these, we modified the 3’ end of the gene by fusing it to a C-terminal HA3 epitope tag followed by a *loxN* site, a T2A peptide and an out-of-frame GFP sequence (Supplementary Figure 5a-b). Attempts to detect and localise the tagged protein in the resulting line (*pficm1:HA*) by western blot and IFA were unsuccessful, possibly due to low abundance of the protein (Supplementary Figure 5c). However, *pficm1:HA* parasites displayed normal replication rates, indicating that the modification had no impact on parasite fitness (Supplementary Figure 5d). In a second gene manipulation step, we replaced intron 1 of *pficm1* in the *pficm1:HA* line with a *2loxPint* element (Koussis et al., 2020), creating line *pficm1:cKO* (Supplementary Figure 5e). DiCre-mediated recombination between the two *loxN* sites (one within the *2loxPint* and one downstream of the HA tag) was predicted to severely truncate the *pficm1* gene while simultaneously positioning the downstream GFP in frame, enabling its expression as a reporter of successful excision (Supplementary Figure 5f-g).

Monitoring of DMSO and RAP-treated *pficm1:cKO* parasites under standard static culture conditions showed complete arrest of growth in the RAP-treated cultures (Fig. 5a). RAP-treated parasites formed morphologically normal mature schizonts at the end of the erythrocytic cycle of treatment (cycle 0), indicating no effects on intracellular development. However, very few schizonts from the RAP-treated cultures underwent egress, resulting in a significant reduction in newly invaded rings (Fig. 5b). To seek further insights we monitored the behaviour of DMSO- and RAP-treated *pficm1:cKO* schizonts by time-lapse video microscopy. As shown in Fig 5c and Supplementary Movie 2, PfICM1-null schizonts exhibited an abnormal phenotype, with either complete loss of egress, or rounding up with increased intracellular merozoite mobility (indicative of parasitophorous vacuole membrane rupture (Thomas et al., 2018)) but no obvious rupture of the RBC membrane, with very few merozoites escaping the confines of the RBC. Quantification of the videos showed fewer egress events and a significant delay in time to egress for the PfICM1-null parasites (Fig. 5d and e). No examples were observed of the explosive egress typical of WT parasites, although following prolonged incubation free PfICM1-null merozoites were visible (Fig. 5b and Supplementary Fig. 6a). To further characterise the phenotype, we compared egress of both DMSO- and RAP-treated *pfpkg:cKO* and *pficm1:cKO* schizonts by monitoring the appearance in culture supernatants of the parasitophorous vacuole protein SERA5. No release of SERA5 ocurred in cultures of PfPKG-null parasites, whilst in marked contrast processed SERA5 appeared in the medium of PfICM1-null parasites, albeit with a delay compared to DMSO-treated parasites (Supplementary Figure 6b). Since proteolytic processing of SERA5 requires discharge of the protease SUB1 from exonemes (Collins et al., 2013b), this result suggests that protein discharge from these secretory organelles is not affected by loss of ICM1. No differences in cellular cGMP levels were detectable in DMSO and RAP treated *pficm1:cKO* parasites (Fig. 5f). Collectively, these results suggested a role for PfICM1 in egress downstream of PKG activation.

**Figure 5.**
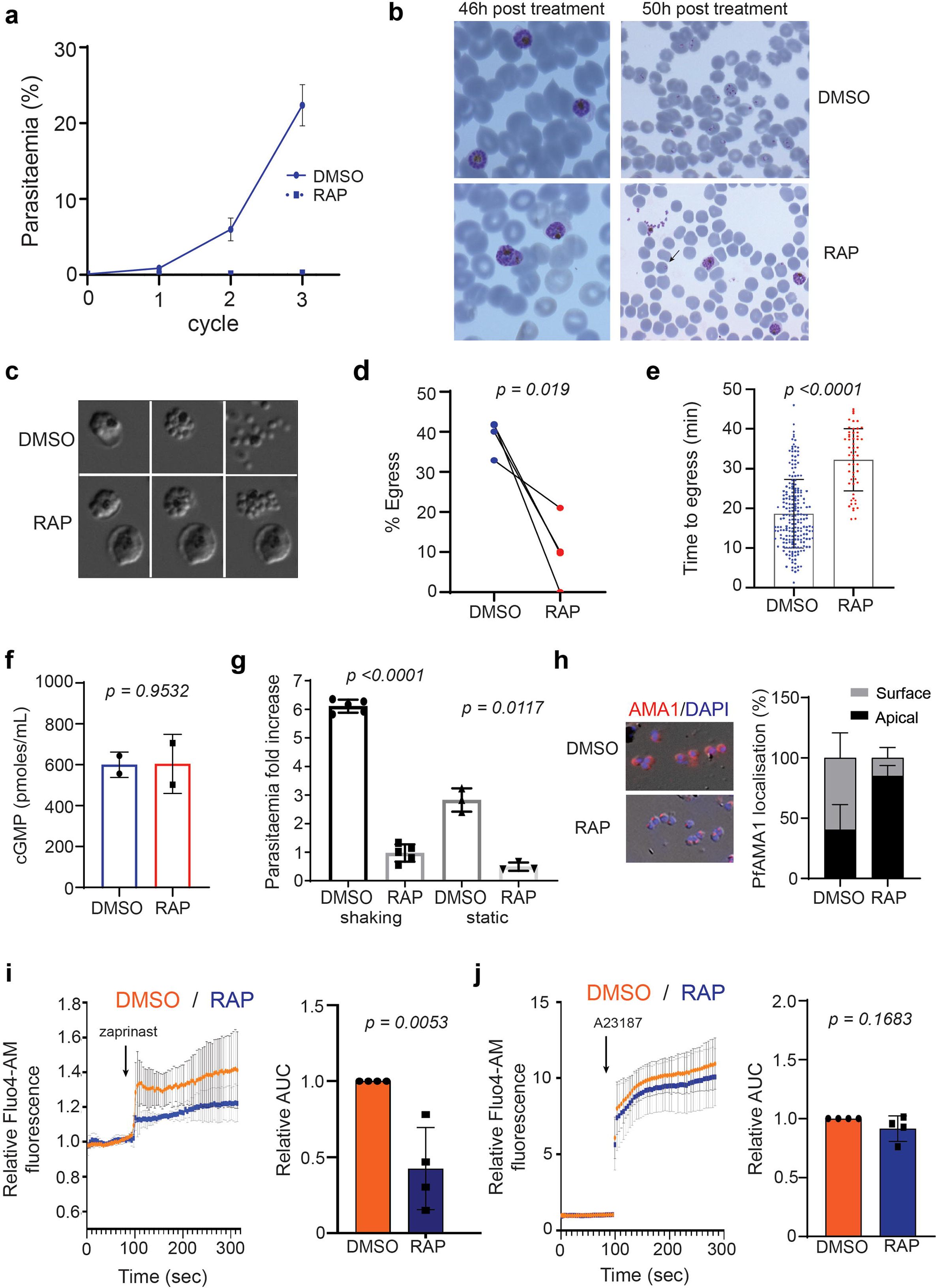
PfICM1 is essential for parasite viability, microneme discharge, efficient egress and invasion. **a)** Replication of DMSO- and RAP-treated *pficm1:cKO* parasites over three erythrocytic cycles. Parasitaemia was measured by flow cytometry. Values are averages of 3 biological replicates. Error bars, ± S.D. **b)** Representative Giemsa-stained blood films showing normal development of DMSO- and RAP-treated *pficm1:cKO* schizonts 46 h post treatment. Over the ensuing 4 h most of DMSO treated parasites successfully invaded as shown by the appearance of newly invaded rings, while RAP-treated parasites displayed a mixed phenotype of very mature schizonts, merozoite clusters and very few newly formed rings (black arrow). **c)** Stills from time-lapse DIC video microscopic monitoring of egress of DMSO and RAP treated *pficm1:cKO* schizonts **d)** Quantification of egress of DMSO and RAP-treated *pficm1:cKO* schizonts based on visual analysis of 45 min-long videos. For each video, both populations were mixed in equal proportions. 4 videos were quantified from 4 independent experiments (two points overlap as they had very similar values). *p*-value was derived from paired *t* tests. **e)** Quantification of the time to egress for each schizont in the videos used for the data in panel **d**. *p*-value was derived from unpaired two tailed *t* tests (DMSO=209 schizonts, RAP=55 schizonts) Error bars, ± S.D. **f)** cGMP levels in DMSO- and RAP-treated *pficm1:cKO* schizonts. Results are from 2 independent experiments, in triplicate. Mean values of each experiment are shown. *p*-value was derived from paired *t* test. Error bars, ± S.D. **g)** Invasion assay under shaking (n=5) and static (n=3) conditions of DMSO- and RAP-treated *pficm1:cKO* schizonts. *p*-values derived from paired *t* test. Error bars, ± S.D. **h) Left panel:** IFA of DMSO- and RAP-treated *pficm1:cKO* parasites. Representative images of translocated AMA1 in DMSO-treated or micronemal AMA1 in RAP-treated merozoites. **Right panel:** Quantification of AMA1 relocalisation by IFA in DMSO- and RAP-treated *pficm1:cKO* schizonts. In approximately 60% of DMSO-treated schizonts, AMA1 was localised to the merozoite surface, compared to only 15% in RAP-treated parasites. 3 biological replicates with at least 150 schizonts in each were quantified. Values, means plus S.D. **i, j)** Relative Fluo4–AM fluorescence response of DMSO- and RAP-treated *pficm1:cKO* schizonts after treatment with zaprinast or A21387. Fluorescence variation with time is presented as means ± S.D. 4 biological replicates were performed. The area under each curve (AUC.) is shown for each stimulation. DMSO values were normalised to 1. Values, mean ± S.D. *p*-values were derived from two tailed *t* tests. See also Figures S5 and S6.

Given the loss of long-term viability in static cultures of PfICM1-null parasites, we addressed whether the egress defect could be overcome by mechanical shear stress. As shown in Fig. 5g, even under shaking conditions production of new rings was severely compromised in RAP-treated *pficm1:cKO* cultures. This prompted us to examine release of other secretory organelle proteins and in particular of AMA1, an essential invasion protein which translocates from micronemes to the surface of daughter merozoites at or just prior to the point of egress (Healer et al., 2002). As shown in Fig. 5h, comparative IFA revealed a severe defect in AMA1 discharge in the PfICM1-null schizonts. It was concluded that this likely explains the loss of invasion in the PfICM1-null parasites.

Release of micronemal contents has been linked to a calcium signalling pathway essential for egress and invasion, involving at least one calcium-dependent protein kinase, PfCDPK5 (Absalon et al., 2018). To address the question of whether loss of ICM1 leads to a defect in calcium mobilisation, we used the PDE inhibitor zaprinast to artificially activate PKG by prematurely increasing cGMP levels, forcing a PKG-dependent increase in cytosolic calcium levels (Brochet et al., 2014). Calcium levels in PfICM1-null parasites exposed to zaprinast were significantly reduced compared to controls, similar to our findings with P_*ama1*_ICM1 gametocytes. In contrast, treatment with the calcium ionophore A23187 had similar effects in both DMSO and RAP-treated *pficmi:cKO* schizonts, suggesting that intracellular calcium levels are unaffected by loss of ICM1 (Fig. 5i and j). Our results indicate that PfICM1 is involved in downstream transduction of calcium signals essential for egress and invasion in asexual blood stages.

### PfICM1 and PfPKG-dependent phosphoproteomes suggest functional overlap in schizonts

Given the physical interaction between ICM1 and PKG in *P. falciparum* schizonts and their roles in calcium mobilisation, we further explored the functional link between the two proteins by determining the impact of ICM1 depletion on the global schizont phosphoproteome. We reasoned that if ICM1 acts downstream of PKG and if its function is regulated directly or indirectly by PKG-dependent phosphorylation, an overlap would be expected between the two phosphoproteomes.

Employing an approach similar to that previously used for PfPKG, we identified 106 phosphosites from 75 proteins that are significantly hypophosphorylated upon disruption of PfICM1 (Fig. 6a and Supplementary Table 3). Proteins involved in cyclic nucleotide dependent signalling cascades were amongst this set, including ACβ, GCα and CDPK1. Motif analysis of the hypophosphorylated phosphopeptides showed a strict preference for lysine in position −3, resembling a minimal PKA/PKG recognition motif (KxxpS/pT) (Fig. 6b). A similar substrate preference was also described in a recent study on the impact of CDPK5 depletion on the schizont phosphoproteome (Blomqvist et al., 2020). GO term enrichment analysis of this set of 75 proteins showed deregulation of cyclic nucleotide biosynthesis, ATPase activity and RNA splicing (GO:0000375), as also observed in the PfPKG phosphoproteome (Fig. 6c and Supplementary Table 3). Comparison of the two phosphoproteome sets identified 73 phosphosites in 67 proteins which are regulated by both PKG and ICM1 in schizonts (Fig. 6d, Supplementary Table 3). A protein interaction network of these 67 proteins using the STRINGApp in Cytoscape (Lopes et al., 2010) revealed a network of 65 nodes with 76 connecting edges (Supplementary Table 3), with 2 main protein clusters of 21 and 6 proteins respectively. Interrogation of the whole network for functional enrichment showed that the spliceosome was the sole significant pathway identified with five proteins clustered together (Fig. 6e), confirming the observation of RNA splicing deregulation from the GO enrichment analysis. The main 21 node cluster incorporated proteins involved in signalling as well as in egress and invasion, including CDPK1, DGK1, ACβ, RHOPH3 and AMA1. Collectively, these data strongly support a functional link between the essential roles of PKG and ICM1.

**Figure 6:**
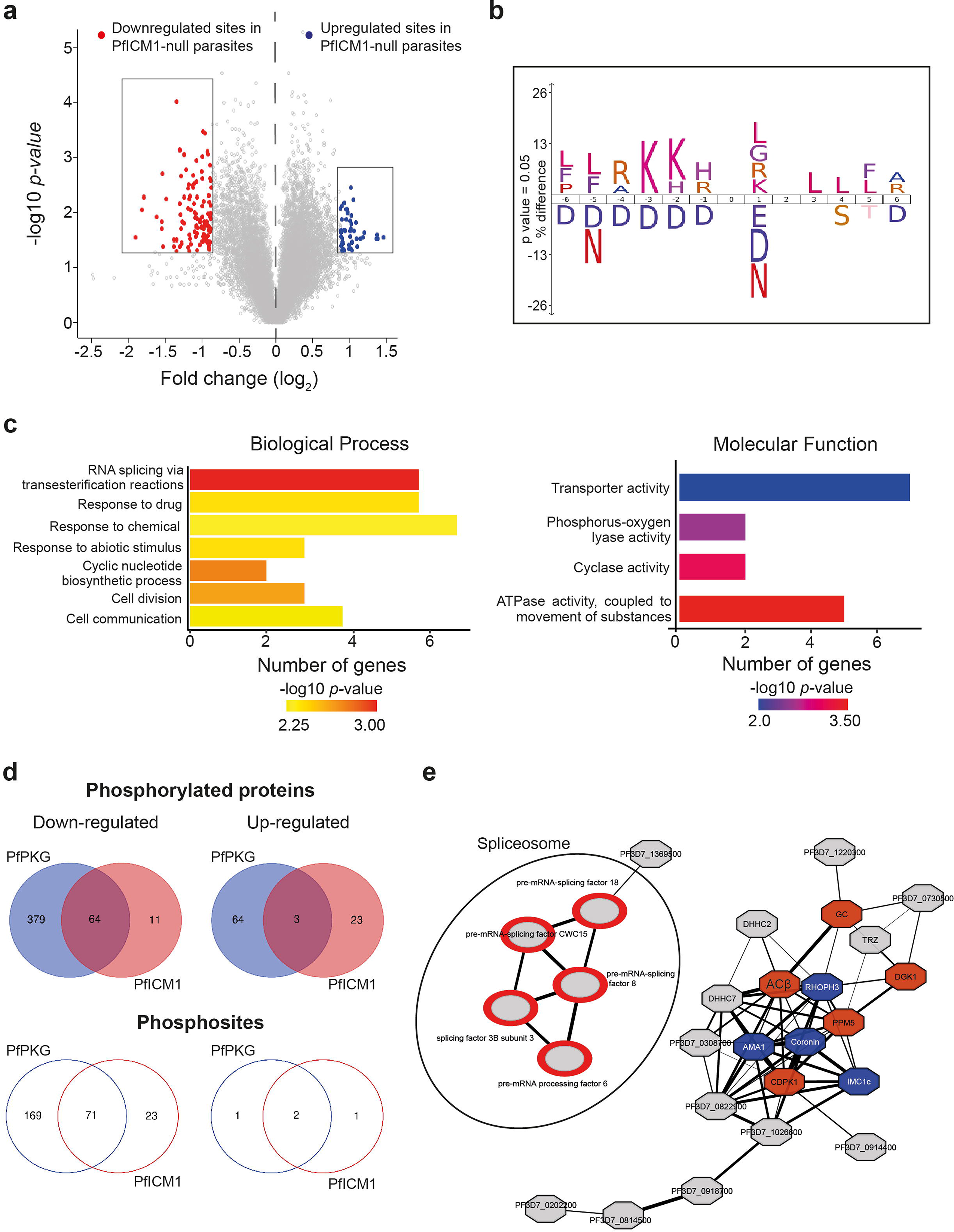
Impact of PfICM1 disruption on the schizont phosphoproteome. **a)** Volcano plot showing distribution of phosphopeptide abundance in the presence or absence of PfICM1. Red circles correspond to significantly down-regulated peptides and blue circles to significantly up-regulated peptides. A cut-off of welch Difference <−0.875 and >0.875 with a p value <0.05 and a localisation probability of >0.7 represents significantly regulated peptides. **b)** Motif analysis of the significantly down-regulated phosphosites in PfICM1-null parasites. All detected phosphopeptides were used as the reference dataset, with amino acids below the position line indicating residues unfavoured for this position. Position 0 corresponds to the phosphorylated residue. **c)** GO enrichment analysis of Biological process and Molecular Function for proteins significantly hypophosphorylated in PfICM1-null parasites. Enriched terms with a p-value <0.01 are shown. **d)** Venn diagram showing the number of phosphoproteins and the corresponding phosphopeptides significantly deregulated by PfPKG and PfICM1. **e)** Interaction-network analyses of common PfPKG/PfICM1 deregulated proteins. The STRING database in Cytoscape was used to create the network with high confidence values (0.7) being used as cut-offs for putative interactions. Gene names for each protein are shown. Proteins involved in signaling pathways are shown as dark-red octagon nodes, while proteins involved in egress and/or invasion as blue octagons. The 5 proteins involved in spliceosome regulation are shown separately as grey nodes with a red doughnut circle. Edge weight corresponds to STRING-defined confidence values, where a thicker line indicates a higher confidence prediction. Conserved proteins with unknown function are shown with their PlasmoDB ID number.

## Discussion

Many genetic and chemical genetic studies have shown a clear requirement for PKG in specific calcium signals or calcium-dependent pathways (Absalon et al., 2018; Brochet et al., 2014; Collins et al., 2013b; Fang et al., 2018; McRobert et al., 2008) but the molecular partners involved in calcium mobilisation have remained elusive (Brochet and Billker, 2016). Here we have identified ICM1 as a tightly-associated PKG partner protein that plays a key role in calcium signals in both schizonts and gametocytes. Remarkably, our data suggest that ICM1 mainly acts downstream of PKG, since ICM1 does not seem to be required for calcium homeostasis during intraerythrocytic development but only for PKG-mediated calcium signals leading to egress and activation of gametogenesis. This suggests a temporally restricted role in calcium homeostasis across the *Plasmodium* life cycle. Future experiments will be needed to determine the impact of ICM1 ablation on CDPK activation in both asexual and sexual stages. Could ICM1 be the long-sought *Plasmodium* IP_3_ receptor, or could it act as a calcium channel? Our modelling data suggest that ICM1 resembles a calcium channel but the low amino acid sequence similarity indicates that it is highly divergent from other well-studied calcium channels. Recombinant and/or heterologous expression and biophysical analysis will be needed to evaluate the function of ICM1 as a calcium channel and to investigate its regulation by IP_3_, other secondary messengers or even by PKG-dependent phosphorylation.

Interactions between kinases and other proteins is a common signalling mechanism, with the A-kinase anchoring proteins (AKAPs) that bind to the cAMP-dependent protein kinase (PKA) or the MAPK scaffolding proteins being some of the best studied examples (Carlisle Michel et al., 2004; Meister et al., 2013). In contrast, little is known about proteins interacting with PKG in eukaryotes as few partners have been identified (Casteel et al., 2002; Schlossmann et al., 2000; Surks et al., 1999; Yuasa et al., 2000). In *Plasmodium*, the only previous data available for a PKG-interacting protein came from an *in vitro* study using the kinase domain of PKG (Govindasamy et al., 2019), whilst a putative interaction between PKG and ICM1 was detected in a very recent elegant study profiling a potent small compound inhibitor of PKG. However, conditional knock down of ICM1 in that work resulted only in a minor growth defect, perhaps due to the different conditional system used (Vanaerschot et al., 2020).

Our identification of the PKG-ICM1 interaction creates new and exciting questions regarding calcium homeostasis in apicomplexan parasites and further strengthens the notion that calcium regulation in this phylum could be regulated by a class of divergent channels.

## Acknowledgements

We thank the excellent service at the bioimaging and flow-cytometry core facilities at the Faculty of Medicine of the University of Geneva. MB is grateful to Oliver Billker for his support during the initial steps of this project. Authors would also like to acknowledge Elena Fernández-Álvaro at GSK for her support in the initial stages of the project and Simon Osborne at LifeArc who was the lead chemist on the project that generated ML10. This work was supported by the Swiss National Science Foundation (starting grant BSSGI0_155852), by the Novartis Foundation (19C189) and by the Fondation privée des Hôpitaux Universitaires de Genève (CONFIRM grant RC05-10) to MB. MB is an INSERM and EMBO young investigator. This work was also supported by Wellcome Trust grant 106239/Z/14/A (KK, AJP and MJB), Wellcome Trust grant 106240/Z/14/Z (DAB), MRC Developmental pathway funding scheme grant (ref G10000779) (DAB) and Wellcome ISSF2 funding to the London School of Hygiene & Tropical Medicine. The work was also supported by funding to MJB from the Francis Crick Institute (https://www.crick.ac.uk/), which receives its core funding from Cancer Research UK (FC001043; https://www.cancerresearchuk.org), the UK Medical Research Council (FC001043; https://www.mrc.ac.uk/), and the Wellcome Trust (FC001043; https://wellcome.ac.uk/).

## Author Contributions

Conceptualization; K.K, D.A.B, M.J.B and M.B. Methodology, Formal Analysis and Investigation; A.C.B, K.K, N.K, S.A.H, H.R.F, M.B, C.P, A.J.P, L.B, P.A, O.S, L.P.B.C, C.W-M, A.H. Writing – Original Draft, A.C.B, K.K, D.A.B, M.J.B and M.B; Writing – Review & Editing; A.C.B, K.K, S.A.H, H.R.F, A.J.P, C,W-M, S.G-D, D.A.B, M.J.B and M.B.; Funding Acquisition; D.A.B, M.J.B and M.B. Resources.; S.G-D A.P.S, D.A.B, M.J.B and M.B. Supervision; S.G-D A.P.S, D.A.B, M.J.B and M.B.

## Declaration of interests

The authors declare no competing interests.

## Methods

### Ethics statement

All animal experiments were conducted with the authorisation numbers GE/82/15 and GE/41/17, according to the guidelines and regulations issued by the Swiss Federal Veterinary Office.

### Reagents

WR99210 was from Jacobus Pharmaceuticals (New Jersey, USA). Rapamycin was from Sigma and used at 20 nM. The PKG inhibitor (4-[7-[(dimethylamino)methyl] −2-(4-fluorphenyl)imidazol[1,2-α]pyridine-3-yl]pyrimidin-2-amine) (compound 2) was stored as a 10 mM stock in DMSO at −20°C, while working dilution was 1 μM. The calcium ionophore A23187 and Zaprinast were both from Sigma and used at 10 μM and 100 μM, respectively.

### *P. falciparum* culture, transfection, synchronisation

All *P. falciparum* lines were maintained at 37 °C in human RBCs in RPMI 1640 containing Albumax II (Thermo-Scientific) and medium was supplemented with 2 mM L-glutamine. Parasite synchronisation and transfections were performed as previously described (Das et al., 2015). Clonal lines were obtained by serial limiting dilution in flat bottomed 96 well plates and single plaques were selected and grown in the presence of 1 μM 5-fluorocytosine (5-FC, provided as clinical grade Ancotyl) to select for Cas9 plasmid-free and marker-free parasites. For parasite genomic DNA extraction (gDNA), the Qiagen DNeasy Blood and Tissue kit was used. Genotype analysis PCR was done using Phusion polymerase (NEB). For *P. falciparum* experiments, DiCre activity was induced by RAP-treatment of early rings (2-3 h post invasion) as previously described (Collins et al., 2013a). Samples for excision PCRs were collected 24 h post RAP treatment.

### *P. berghei* maintenance, purification and transfection

*P. berghei* maintenance, purification and transfection was performed as previously described (Balestra et al., 2020). *P. berghei* ANKA strain (Sebastian et al., 2012) derived clones 2.34 (Moon et al., 2009), and 615 (Philip and Waters, 2015), together with derived transgenic lines, were grown and maintained in CD1 outbred mice. Six to ten week-old mice were obtained from Charles River laboratories, and females were used for all experiments. Mice were specific pathogen free (including *Mycoplasma pulmonis*) and subjected to regular pathogen monitoring by sentinel screening. They were housed in individually ventilated cages furnished with a cardboard mouse house and Nestlet, maintained at 21 ± 2 °C under a 12 h light/dark cycle, and given commercially prepared autoclaved dry rodent diet and water *ad libitum*. The parasitaemia of infected animals was determined by microscopy of methanol-fixed Giemsa-stained thin blood smears.

For gametocyte production, parasites were grown in mice that had been phenyl hydrazine-treated three days before infection. One day after infection, sulfadiazine (20 mg/L) was added in the drinking water to eliminate asexually replicating parasites. Microgametocyte exflagellation was measured three or four days after infection by adding 4 μl of blood from a superficial tail vein to 70 μl exflagellation medium (RPMI 1640 containing 25 mM HEPES, 4 mM sodium bicarbonate, 5% fetal calf serum (FCS), 100 μM xanthurenic acid, pH 7.4). To calculate the number of exflagellation centres per 100 microgametocytes, the percentage of RBCs infected with microgametocytes was assessed on Giemsa-stained smears. For gametocyte purification, parasites were harvested in suspended animation medium (SA; RPMI 1640 containing 25 mM HEPES, 5% FCS, 4 mM sodium bicarbonate, pH 7.20) and separated from uninfected RBCs on a Histodenz (Sigma) cushion made from 48% of a Histodenz stock (27.6% [w/v] Histodenz in 5.0 mM TrisHCl, 3.0 mM KCl, 0.3 mM EDTA, pH 7.20) and 52% SA, final pH 7.2. Gametocytes were harvested from the interface. To induce ICM1-AID/HA degradation, 1 mM auxin dissolved in ethanol was added to purified gametocytes for 1 h.

Schizonts for transfection were purified from overnight *in vitro* culture on a Histodenz cushion made from 55% of the Histodenz stock and 45% PBS. Parasites were harvested from the interface and collected by centrifugation at 500 *g* for 3 min, resuspended in 25 μl Amaxa Basic Parasite Nucleofector solution (Lonza) and added to 10-20 μg DNA dissolved in 10 μl H_2_O. Cells were electroporated using the FI-115 program of the Amaxa Nucleofector 4D. Transfected parasites were resuspended in 200 μl fresh RBCs and injected intraperitoneally into mice. Parasite selection with 0.07 mg/mL pyrimethamine (Sigma) in the drinking water (pH ~4.5) was initiated one day after infection.

### Plasmid construction and genotyping of *P. falciparum* transgenic lines

Oligonucleotides used in this study are shown in Supplementary Table 4.

To create line PKG-GFP, the triple HA tag from plasmid pHH1-PfPKG-HA (Hopp et al., 2012) was replaced by eGFP. In brief, eGFP was PCR amplified from plasmid pBAT-SIL6 (Kooij et al., 2012) using primers GFP_For and GFP_Rev. The PCR product was digested with AvrII and XhoI and ligated to vector pHH1-PfPKG-HA previously digested with the same enzymes. Plasmid was transfected into 3D7 *P. falciparum* and growth medium replaced◻24◻h post transfection with fresh medium containing 2.5◻nM WR99210. Parasites were subjected to culture in the absence of WR99210 (3 cycles) followed by culturing with drug to select for transgenic parasites. Clonal lines were confirmed by PCR using primers wtpkg_For and 3utr_Rev, and maintained in the presence of WR99210.

The DiCre-expressing B11 line (Perrin et al., 2018) was used to generate line *icm:cKO* in two steps. The first step aimed at modifying the 3’ end of the gene. To achieve that, a commercially obtained construct (GeneArt, Thermo) contained in tandem: 1) a 5’ homology arm of 456 bp endogenous and 255 bp of synthetic sequence comprising the 3’ end of exon 2 and the whole of exon 3, 2) a fragment comprising a triple HA tag, *loxN* and the 3’ 46 bp of the *sera2* intron and finally 3) the first 408 bp of *pficm1* 3’ UTR as a 3’ homology arm. This plasmid (pMX_icm1-int) was digested with SalI/SacI and a T2A peptide followed by a GFP gene (fragment obtained from plasmid pT2A_D1_cKO, a kind gift of Dr. Edgar Deu) was cloned in after the 3’ end of the *sera2* intron, resulting in plasmid pMX_icm1_2. A single guide RNA (sgRNA) targeting sequence TTGATTAAATAAATATATAA was inserted into a previously described pDC2 plasmid expressing Cas9, resulting in plasmid pDC2-icm1_2g. The repair plasmid (pMX_icm1_2) was linearised overnight with BgIII and transfected in B11 parasites together with plasmid pDC2_icm1_2g. Integration was confirmed by PCR, using primer pairs L2_For/L2_Rev and L2_For/L2_HA. This clonal line (*icm1:HA*) was obtained via limiting dilution and was used for the 2nd modification step, which involved replacing the endogenous intron 1 with the 2loxPint module (Koussis et al., 2020). A commercially obtained construct (GeneArt, Thermofisher) was used (pMX_icm1_2loxPint), containing the 2loxPint module flanked by approximately 400 bp of endogenous sequence as homology arms. This vector was linearised with BglII and cotransfected with a pDC2 based Cas9 plasmid(Knuepfer et al., 2017), harboring the sgRNA targeting sequence ATTTCAAAGGACTTACTTTA, in line *icm1_HA*. Clonal lines were obtained as described above and integration was confirmed using primer pairs L1_For/L1_Rev.

To create a conditional knockout of PfPKG, vector pDC*_*loxnPKG:lox22mCherry^4^ was digested with XbaI and NheI. The latter restriction site was blunted and the vector was re-ligated resulting in vector pDC*_*lox22mCherry. This plasmid was linearised overnight with ScaI and transfected together with the pDC2-pkg Cas9 plasmid into *P. falciparum* line *pfpkg_2lox* (Koussis et al., 2020). Correct 5’ and 3’ integration were verified by PCR using primers wtpkg_For/5int_Rev. Absence of the endogenous locus was confirmed using primers exon1_For/PKGutr_Rev.

### Plasmid construction and genotyping of *P. berghei* transgenic lines

The oligonucleotides used to generate and genotype the mutant parasite lines are in Supplementary Table 4.

#### Restriction/ligation cloning

To generate the P_*ama1*_ICM1 line, the plasmid pOB116-ama1 was used (Sebastian et al., 2012). The first 1000 bp from *PbICM1* gene was amplified from genomic DNA using the primers 453 and 458. The PCR product was digested with XhoI and NotI and ligated into the pOB116-ama1 plasmid previously digested with the same enzymes, resulting in the plasmid pAB025. The last 750 bp of *PbICM1* 5’ UTR were amplified using primers 459 and 464 and the resulting fragment was digested with HindIII and PstI, and ligated into pAB026 previously digested with the same enzymes. The same strategy was used to generate the line P_*clag9*_ICM1, with the plasmid pOB116-clag9 resulting in plasmid pAB027. Both plasmids were digested with the enzymes EcoRV and HindIII to be transfected in *P. berghei.*

#### Recombineering

ICM1-HA3 and ICM1-AID/HA tagging constructs were generated using phage recombineering in *Escherichia coli* TSA strain with PlasmoGEM vectors (http://plasmogem.sanger.ac.uk/) as previously described (Pfander et al., 2013; Pfander et al., 2011). The modified library inserts were then released from the plasmid backbone using NotI. The ICM1-AID/HA (PbGEM-645751) targeting vector was transfected into the 615 line (Philip and Waters, 2015). The ICM1-HA3 (PbGEM-121338) and ICM1-KO (PbGEM-265644) vectors were transfected into the 2.34 line.

Each mutant parasite was genotyped by PCR using three combinations of primers, specific for either the WT or the modified locus on both sides of the targeted region (experimental designs are shown in Supplemental Figures). Controls using wild type DNA were included in each genotyping experiment; parasite lines were cloned when indicated.

### *P. falciparum* growth assays

Growth rates in *P. falciparum* mutant lines were performed as follows: synchronous ring-stage parasites at 0.1% parasitaemia and 2% haematocrit were dispensed in triplicate into 12-well plates. 50 μl samples from each well were collected at 0, 2, 4 and 6 days, stained with SYBR green and analysed by flow cytometry on a BD FACSVerse™ using BD FACSuite™ software. Data acquired were analysed using FlowJo software.

### *P. falciparum* egress and invasion assays

Mature schizonts were isolated by Percoll centrifugation and incubated for a further 3 h in complete medium containing the reversible PKG inhibitor compound 2 (1 μM). After removal of the inhibitor, schizonts were immediately resuspended in fresh serum-free RPMI at 37°C to allow egress. Schizont pellets and culture supernatants at t=0 were collected as a control sample, whilst culture supernatants were collected by centrifugation at the relevant time points.

For invasion assays, mature schizonts were incubated at 2-4% parasitaemia in 2% haematocrit under static or shaking conditions for 4h. Unruptured schizonts were removed by Percoll gradient, parasitaemia was measured again and the fold-increase was calculated.

### Calcium measurements

#### *P. falciparum* schizonts

Changes in calcium levels were measured using Fluo-4, AM (Thermo) loaded parasites and differences in fluorescence were measured over time on a BD FACSVerse™ flow cytometer using BD FACSuite™ software. In brief, schizonts were washed twice in phenol red- and serum-free RPMI medium (11835030, Thermo). 5 μM Fluo-4, AM was added to 1◻ml of parasite preparation. Cells were incubated with Fluo-4, AM at 37◻°C for 45◻min and washed twice in phenol red- and serum free RPMI as above. Parasites were incubated for 30◻min for de-esterification followed by a further two washes. Schizonts were resuspended in 1 ml of warm phenol red- and serum free RPMI. For Fluo-4, AM detection a 488 nm Blue Laser was used with a 530/30 filter. Fluorescence levels were measured for a period of 90 sec to obtain a baseline read. Compounds were then added onto the cells and reads recorded for a further 3 min. Data analysis was done with FlowJo software.

#### *P. berghei* gametocytes

To measure changes in calcium level, *P. berghei* gametocytes were harvested in coelenterazine loading buffer(Billker et al., 2004) (CLB; PBS containing 20 mM HEPES, 20 mM Glucose, 4 mM NaHCO3, 1 mM EDTA, 0.1% BSA, pH: 7.2). Gametocytes were purified as described above and resuspended in 1 ml of CLB and loaded with 5 μM of Fluo4-AM for an hour. The parasites were then washed twice with CLB, resuspended in 500 μl and 20 μl were added per well in a 96 well plate. Compound dispense and simultaneous fluorescence reading were performed by a FDSS μCELL (Hamamatsu Photonics) for a period of 5 min.

### Flow cytometry analysis of gametocytes

To analyse their DNA content, gametocytes were harvested in SA and purified as described above. Gametocytes were resuspended in 100 μl and activated with 100 μl of modified exflagellation medium (RPMI 1640 containing 25 mM HEPES, 4 mM sodium bicarbonate, 5% fetal calf serum, 200 μM xanthurenic acid, pH 8.0). Gametogenesis was stopped by ice-cold PBS containing Vybrant dye cycle violet (Life technologies) and cells were stained for 30 min at 4◻C. DNA content was determined using a Beckman Coulter Gallios 4 and analysed with the Kaluza software. Per sample, >10,000 cells were analysed.

### Immunoblotting and Immunofluorescence assays

#### P. falciparum

Synchronised *P. falciparum* schizonts were isolated by Percoll gradient centrifugation and washed in RPMI 1640 without Albumax. Parasites were extracted into a Triton X-100 buffer (20 mM Tris-HCl pH 7.4, 150 mM NaCl, 0.5 mM EDTA, 1% Triton X-100, supplemented with 1× protease inhibitors - Roche). Extracts were incubated on ice for 30 min then clarified by centrifugation at 12,000g for 15 min at 4°C. Supernatants were mixed with SDS sample buffer containing DTT and incubated for 5 min at 95°C prior to fractionation by SDS-PAGE analysis on 4%–15% Mini-PROTEAN TGX Stain-Free Protein Gels (Bio-Rad). Transfer to nitrocellulose membranes and probing for Western blot analysis was as described previously. A rabbit polyclonal human-PKG antibody (Enzo) was used for PKG detection at 1:1000. A GFP-specific monoclonal antibody (mAb) 11814460001 (Roche) was used at 1:1000, as was a polyclonal rabbit anti-mCherry (ab167453; Abcam). A polyclonal rabbit anti-SERA5 antibody was used at 1:2000 and an anti-HSP70 abtibody (kind gift of Dr Ellen Knuepfer, Francis Crick Institute) was used at 1:2000.

For IFA analysis, thin blood smears of *pficm1:HA* percoll enriched schizonts were fixed in 4% (w/v) paraformaldehyde and permeabilised in 0.1% (v/v) Triton X-100. After blocking with 3% BSA, smears were probed with a rat anti-HA (3F10), 1:100 and a mouse MSP-1 (1:1000). For AMA1 relocalisation, vehicle (DMSO) or RAP-treated Percoll-enriched *pficm1:cKO* schizonts were probed with a rabbit anti-AMA1, 1:200. Subsequently, slides were probed with the required Alexa Fluor 594-conjugated secondary antibody diluted 1:1,000. Images were collected using a Nikon Eclipse Ni-E wide field microscope with a Hamamatsu C11440 digital camera and 100x/1.45NA oil immersion objective. Identical exposure conditions were used at each wavelength for each pair of mock- and RAP-treated samples under comparison. Images were processed using Fiji software (Schindelin et al., 2012).

#### P. berghei

Gametocyte immunofluorescence assays were performed as previously described(Volkmann et al., 2012). For HA and α-tubulin staining, purified cells were fixed with 4% paraformaldehyde and 0.05% glutaraldehyde in PBS for 1 h, permeabilised with 0.1% Triton X-100/PBS for 10 min and blocked with 2% BSA/PBS for 2 h. Primary antibodies were diluted in blocking solution (rat anti-HA clone 3F10, 1:1000; mouse anti-α-tubulin clone DM1A, 1:1000, both from Sigma-Aldrich). Anti-rat Alexa594, anti-mouse Alexa488, anti-rabbit Alexa 488, Anti-rabbit Alexa594 were used as secondary antibodies together with DAPI (all from Life technologies), all diluted 1:1000 in blocking solution. Confocal images were acquired with a LSM700 or a LSM800 scanning confocal microscope (Zeiss).

### Time-lapse and live fluorescence microscopy

Viewing chambers were constructed as previously described (Collins et al., 2013b). Images were recorded on a Nikon Eclipse Ni light microscope fitted with a Hamamatsu C11440 digital camera and Nikon N Plan Apo λ 63x/1.45NA oil immersion objective. For time-lapse video microscopy, vehicle (DMSO) or RAP-treated schizonts were synchronised by incubation with the reversible PKG inhibitor compound 2, then washed, mixed in equal proportions and monitored for egress. Differential inference contrast (DIC) images were taken at 10 sec intervals over 45 min while fluorescence (GFP or mCherry) images were taken every 5-10 min to prevent bleaching. Time-lapse videos were analysed and annotated using Fiji (Schindelin et al., 2012).

### Measurement of intracellular cyclic nucleotide levels

Relative intracellular cGMP levels in mature schizonts were measured using the Direct cGMP ELISA kit (Enzo). RAP- or DMSO-treated *icm1:cKO* schizonts were Percoll purified and lysed in the presence of 0.1 ml of 0.1 M HCl to harvest cGMP. Samples were incubated at room temperature for 5 minutes with intermittent vortexing. Standards were fitted to a sigmoidal curve and used to determine cyclic nucleotide concentrations in parasite samples.

### ICM1 protein structure prediction

The homology models of PfICM1 were constructed initially using the Phyre2 Server in intensive mode (Kelley et al., 2015). The I-TASSER meta-threading procedure (Iterative Threading ASSEmbly Refinement; https://zhanglab.ccmb.med.umich.edu/I-TASSER/) was run to generate a molecular model of PfICM1, using the first 1,500 amino acids of the protein (I-TASSER maximum allowed residues) and covering domains of the protein predicted to be highly helical (Roy et al., 2010; Yang et al., 2015; Zhang, 2008). The local Meta-Threader Server identified PDB 6MU1, 6JXC, 6RLB, 6C08 as the top four threading templates for protein structure prediction, three of them being transporter proteins.

### Immunoprecipitation and mass-spectrometry analysis in *P. falciparum*

For pull-down analysis, frozen schizont preparations from lines PfPKG-HA, PKG-GFP and 3D7 were resuspended in 100 μl lysis buffer (20 mM Tris-HCl pH 7.5, 150 mM NaCl, 1 mM EDTA, 1 % TX-100, complete protease inhibitors and PhosSTOP) and extracted at 4 °C for 30 min with intermittent vortexing. Extracts were clarified by centrifugation at 16,000 × g for 20 min at 4 °C, filtered through 0.22 Corning® Costar® Spin-X® centrifuge tube filters, and then incubated with 25 μl anti-HA magnetic beads (Pierce) or with 25 μL GFP-Trap Magnetic Agarose (Chromotek) overnight at 4 °C. Sample processing followed the manufacturer’s protocol. Protein complexes were eluted from beads with the addition of 2x volumes of reduced 2× SDS sample buffer. Samples were incubated at 95◻°C for 5 min and run on 12% SDS–PAGE gels. Band slices covering the whole separation profile were excised and washed. Reduced and alkylated proteins were in-gel digested with 100 ng trypsin (modified sequencing grade, Promega) overnight at 37°C. Supernatants were dried in a vacuum centrifuge and resuspended in 0.1% TriFluoroAcetic acid (TFA). 1-10 μL of digested protein acidified to a final concentration of 0.1% TFA was loaded on an Ultimate 3000 nanoRSLC HPLC (Thermo Scientific) at 15 μl/min of 0.1% TFA onto a 2 mm × 0.3 mm Acclaim Pepmap C18 trap column (Thermo Scientific) prior to the trap being switched to elute at 0.25 μl/min through a 50cm × 75 μm EasySpray C18 column. A 90’ gradient of 9%−25% B over 37’, then 25%-40% B over 18’ was used followed by a short gradient to 100% B and back down to 9% B and a 20’ equilibration in 9% B (A= 2%ACN,0.1% formic acid; B= 80%ACN, 0.1% formic acid).

The Orbitrap was operated in “Data Dependent Acquisition” mode with a survey scan at a resolution of 120,000 from m/z 300-1500, followed by MS/MS in “TopS” mode. Dynamic exclusion was used with a time window of 20 sec. The Orbitrap charge capacity was set to a maximum of 1e6 ions in 10 ms, whilst the LTQ was set to 1e4 ions in 100 ms. Raw files were processed using Maxquant 1.3.0.5 and Perseus 1.4.0.11 against a recent version of PlasmoDB (www.plasmodb.org). A decoy database of reversed sequences was used to filter false positives, at a peptide false detection rate of 1%.

### Immunoprecipitation and mass-spectrometry analysis in *P. berghei*

#### Sample preparation

Co-immunoprecipitations (IPs) of proteins were performed with purified gametocytes. The following IPs were performed: ICM1-HA3 in schizonts or gametocytes (15 sec pa +/− Compound A), PKG-HA3 in schizonts or gametocytes (15 sec pa +/− Compound A). IPs from wild type non-activated gametocytes lacking an epitope tag were used as controls.

Samples were fixed for 10 min with 1% formaldehyde. Parasites were lysed in RIPA buffer (50 mM Tris HCl pH 8, 150 mM NaCl, 1% NP-40, 0.5% sodium deoxycholate, 0.1% SDS) and the supernatant was subjected to affinity purification with anti-HA antibody (Sigma) conjugated to magnetics beads. Beads were re-suspended in 100 μl of 6 M urea in 50 mM ammonium bicarbonate (AB). Two μl of 50 mM dithioerythritol (DTE) were added and the reduction was carried out at 37°C for 1h. Alkylation was performed by adding 2 μl of 400 mM iodoacetamide for 1 h at room temperature in the dark. Urea was reduced to 1 M by addition of 500 μl AB and overnight digestion was performed at 37 °C with 5 μl of freshly prepared 0.2 μg/μl trypsin (Promega) in AB. Supernatants were collected and completely dried under speed-vacuum. Samples were then desalted with a C18 microspin column (Harvard Apparatus) according to manufacturer’s instructions, completely dried under speed-vacuum and stored at −20°C.

#### Liquid chromatography electrospray ionisation tandem mass spectrometry (LC-ESI-MSMS)

Samples were diluted in 20 μl loading buffer (5% acetonitrile [CH_3_CN], 0.1% formic acid [FA]) and 2 μl were injected onto the column. LC-ESI-MS/MS was performed either on a Q-Exactive Plus Hybrid Quadrupole-Orbitrap Mass Spectrometer (Thermo Fisher Scientific) equipped with an Easy nLC 1000 liquid chromatography system (Thermo Fisher Scientific) or an Orbitrap Fusion Lumos Tribrid mass Spectrometer (Thermo Fisher Scientific) equipped with an Easy nLC 1200 liquid chromatography system (Thermo Fisher Scientific). Peptides were trapped on an Acclaim pepmap100, 3 μm C18, 75 μm × 20 mm nano trap-column (Thermo Fisher Scientific) and separated on a 75 μm × 250 mm (Q-Exactive) or 500 mm (Orbitrap Fusion Lumos), 2 μm C18, 100 Å Easy-Spray column (Thermo Fisher Scientific). The analytical separation used a gradient of H_2_O/0.1% FA (solvent A) and CH_3_CN/0.1 % FA (solvent B). The gradient was run as follows: 0 to 5 min 95 % A and 5 % B, then to 65 % A and 35 % B for 60 min, then to 10 % A and 90 % B for 10 min and finally for 15 min at 10 % A and 90 % B. Flow rate was 250 nL/min for a total run time of 90 min.

Data-dependant analysis (DDA) was performed on the Q-Exactive Plus with MS1 full scan at a resolution of 70,000 Full width at half maximum (FWHM) followed by MS2 scans on up to 15 selected precursors. MS1 was performed with an AGC target of 3 × 10^6^, a maximum injection time of 100 ms and a scan range from 400 to 2000 m/z. MS2 was performed at a resolution of 17,500 FWHM with an automatic gain control (AGC) target at 1 × 10^5^ and a maximum injection time of 50 ms. Isolation window was set at 1.6 m/z and 27% normalised collision energy was used for higher-energy collisional dissociation (HCD). DDA was performed on the Orbitrap Fusion Lumos with MS1 full scan at a resolution of 120,000 FWHM followed by as many subsequent MS2 scans on selected precursors as possible within a 3 sec maximum cycle time. MS1 was performed in the Orbitrap with an AGC target of 4 × 10^5^, a maximum injection time of 50 ms and a scan range from 400 to 2000 m/z. MS2 was performed in the Ion Trap with a rapid scan rate, an AGC target of 1 × 10^4^ and a maximum injection time of 35 ms. Isolation window was set at 1.2 m/z and 30% normalised collision energy was used for HCD.

#### Database searches

Peak lists (MGF file format) were generated from raw data using the MS Convert conversion tool from ProteoWizard. The peak list files were searched against the PlasmoDB_*P.berghei* ANKA database (PlasmoDB.org, release 38, 5076 entries) combined with an in-house database of common contaminants using Mascot (Matrix Science, London, UK; version 2.5.1). Trypsin was selected as the enzyme, with one potential missed cleavage. Precursor ion tolerance was set to 10 ppm and fragment ion tolerance to 0.02 Da for Q-Exactive Plus data and to 0.6 for Lumos data. Variable amino acid modifications were oxidized methionine and deamination (Asn and Gln) as well as phosphorylated serine, threonine and tyrosine. Fixed amino acid modification was carbamidomethyl cysteine. The Mascot search was validated using Scaffold 4.8.4 (Proteome Software) with 1% of protein false discovery rate (FDR) and at least 2 unique peptides per protein with a 0.1% peptide FDR.

#### PCA analysis

Enrichment and principal component analysis were performed in the statistical programming package ‘R’ (www.r-project.org). Quantitative values were analysed as log-transformed spectral count values and displayed in principal components with greatest degrees of variance.

### Chemoproteomics

Kinobeads were prepared as described (Bantscheff et al., 2011; Bergamini et al., 2012). The chemoproteomic affinity capturing experiments were performed as previously described (Bantscheff et al., 2011; Matralis et al., 2019). Briefly, Kinobeads were incubated with *P. falciparum* extract (Paquet et al., 2017), which was pre-incubated with compound or DMSO (vehicle control). The experimental set up was such that 10 samples are measured in parallel (TMT 10-plex (Werner et al., 2014)) to generate values for the affinity of the beads to the bound proteins (“depletion” values, 4 samples) and to generate IC_50_ values based on dose-dependent competition (6 samples, 20 μM, 1:3 dilutions) in a single experiment. Beads were transferred to Filter plates (Durapore PVDF membrane, Merck Millipore), washed extensively and eluted with SDS sample buffer (Eberl et al., 2019).

Proteins were digested according to a modified single pot solid-phase sample preparation (SP3) protocol (Hughes et al., 2014; Moggridge et al., 2018). Proteins were digested by resuspending in 0.1 mM HEPES (pH 8.5) containing TCEP, Chloracetamide, Trypsin and LysC following o/n incubation.

Peptides were labeled with isobaric mass tags (TMT10, Thermo FisherScientific, Waltham, MA using the 10-plex TMT reagents, enabling relative quantification of 10 conditions in a single experiment (Werner et al., 2012; Werner et al., 2014). Labeled peptide extracts were combined to a single sample per experiment, lyophilized and subjected to LC-MS analysis.

Samples were injected into an Ultimate3000 nanoRLSC (Dionex) coupled to a Q-Exactive mass spectrometer (Thermo Fisher Scientific). Peptides were separated on reversed-phase columns (Reprosil) at 55°C g by gradient elution (3.5 % acetonitrile to 29 % acetonitrile in aqueous 0.1% formic acid, 3.5 % DMSO) in 120 minutes. The instruments were operated with Tune 2.4 and Xcalibur 3.0 build 63.

Mascot 2.4 (Matrix Science, Boston, MA) was used for protein identification by using a 10 parts per million mass tolerance for peptide precursors and 20 mD (HCD) mass tolerance for fragment ions. To create the fasta file for mascot searching, all proteins corresponding to the taxonomy ‘Plasmodium falciparum (isolate 3D7)’ were downloaded from Uniprot (release 20170621) and supplemented with common contaminant protein sequences of bovine serum albumin, porcine trypsin and mouse, rat, sheep and dog keratins. To assess the false discovery rate (FDR) “decoy” proteins (reverse of all protein sequences) were created and added to the database, resulting in a database containing a total of 14266 protein sequences, 50% forward, 50% reverse.

Unless stated otherwise, we accepted protein identifications as follows: (i) For single-spectrum to sequence assignments, we required this assignment to be the best match and a minimum Mascot score of 31 and a 10× difference of this assignment over the next best assignment. Based on these criteria, the decoy search results indicated <1% false discovery rate (FDR). (ii) For multiple spectrum to sequence assignments and using the same parameters, the decoy search results indicated <0.1% FDR. Quantified proteins were required to contain at least 2 unique peptide matches. FDR for quantified proteins was < 0.1%.

### *P. falciparum* phosphoproteomics

Parasite pellets were treated as previously described (Patel et al., 2019). In brief, schizont pellets were resuspended in 1 mL 8M urea in 50 mM HEPES, pH 8.5, containing protease and phosphatase inhibitors and 100 U/mL benzonase (Sigma). Proteins were extracted from the pellets by sonication. Samples were incubated on ice for 10 min and centrifuged for 30 min at 14,000 rpm at 4°C. Protein content was estimated by a BCA protein assay and 200 μg of each sample were reduced with 10 mM dithiothreitol for 25 min at 56°C and then alkylated with 20 mM iodoacetamide for 30 min at room temperature. Samples were digested initially with LysC (WAKO) at 37°C for 2.5 h and subsequently with 10 μg trypsin (modified sequencing grade, Promega) overnight. After acidification, C_18_ MacroSpin columns (Nest Group) were used to clean up the digested peptide solutions and the eluted peptides dried by vacuum centrifugation. Samples were resuspended in 50 mM HEPES and labelled using the 0.8 mg Tandem Mass Tag 10plex isobaric reagent kit (Thermo Scientific) following manufacturer’s instructions. The eluted TMT-labelled peptides were dried by vacuum centrifugation and phosphopeptide enrichment was subsequently carried out using the sequential metal oxide affinity chromatography (SMOAC) strategy with High Select TiO_2_ and Fe-NTA enrichment kits (Thermo Scientific). Eluates were combined prior to fractionation with the Pierce High pH Reversed-Phase Peptide Fractionation kit (Thermo Scientific). The dried TMT-labelled phosphopeptide fractions generated were resuspended in 0.1% TFA for LC-MS/MS analysis using a U3000 RSLCnano system (Thermo Scientific) interfaced with an Orbitrap Fusion Lumos (Thermo Scientific). Each peptide fraction was pre-concentrated on an Acclaim PepMap 100 trapping column before separation on a 50-cm, 75-μm I.D. EASY-Spray Pepmap column over a three hours gradient run at 40°C, eluted directly into the mass spectrometer. The instrument was run in data-dependent acquisition mode with the most abundant peptides selected for MS/MS fragmentation. Two replicate injections were made for each fraction with different fragmentation methods based on the MS^2^ HCD and MSA SPS MS^3^ strategies described (Jiang et al., 2017). The acquired raw mass spectrometric data were processed in MaxQuant (Cox and Mann, 2008) (v 1.6.2.10) and searched against UniProt *Homo Sapiens* complete proteome canonical sequences (September 2019) and PlasmoDB. Fixed modifications were set as Carbamidomethyl (C) and variable modifications set as Oxidation (M) and Phospho (STY). The estimated false discovery rate was set to 1% at the peptide, protein, and site levels. A maximum of two missed cleavages were allowed. Reporter ion MS^2^ or Reporter ion MS^3^ was appropriately selected for each raw file. Other parameters were used as preset in the software. The MaxQuant output file PhosphoSTY Sites.txt, an FDR-controlled site-based table compiled by MaxQuant from the relevant information about the identified peptides, was imported into Perseus (v1.4.0.2) for data evaluation. Sequence logos were generated by IceLogo (https://iomics.ugent.be/icelogoserver/). Paired t-tests were performed using R (R Core Team, 2017; https://www.R-project.org/) Significantly down-regulated genes from PfPKG and PfICM1 phosphoproteomes were used for enrichment of GO terms in PlasmoDB using a cut-off p-value of 0.05. Results were visualised using REVIGO (Supek et al., 2011) with the *Plasmodium falciparum* database and similarity: 0.7, semantic similarity measure: SimRel. Plots were made using R (R Core Team, 2017; https://www.R-project.org/) package ggplot2(Wickham, 2016). Common deregulated proteins were imported to Cytoscape v. 3.8.0 via the STRING database App using a confidence threshold > 0.7 (Doncheva et al., 2019; Lopes et al., 2010; Szklarczyk et al., 2019).

### *P. berghei* phosphoproteomics

#### Sample preparation

Samples (about 100 μl of packed cell volume) were diluted with 25 μl of Laemmli 5x buffer, and sonicated 6 × 30 sec at 70% amplitude and 0.5 pulse (30 sec. on ice between each cycle). Samples were heated 5 min at 65°C and loaded onto a Novex Wedgewell 12% Tris-glycine gel (Invitrogen). Each sample was loaded on two different wells and protein’s migration was then performed according to manufacturer’s instructions. Proteins were then stained with coomassie blue. For each lane, a band between 180 kDa and 220 kDa was cut and destained by incubation in 100 μl of 30% acetonitrile (AcN) in 50 mM ammonium bicarbonate (AB) for 15 min at room temperature. Proteins were reduced by incubation of gel pieces for 35 min. at 56°C in 100 μl of 10 mM DTE in 50 mM AB. DTE solution was then replaced by 100 μl of 55 mM iodoacetamide in 50 mM AB and proteins were alkylated by incubation of the gel pieces for 30 min at room temperature in the dark. Gel pieces were then washed for 10 min with 100 μl of 50 mM AB and dehydrated for 10 min with 100 μl of 30% AcN. Gel pieces were then dried for 30 min in a vacuum centrifuge. Dried pieces of gel were rehydrated for 45 min at 4°C with 70 μl of trypsin (6.25 ng/μl in AB 50 mM) and then incubated overnight at 37°C. Supernatant was transferred to a new polypropylene tube and two additional peptide extraction were performed with respectively 70 μl of 1% TFA and 100 μl of 50% ACN; 0.1% TFA. Extractions were pooled and dried under speed vacuum.

#### Titanium dioxide enrichment

Phosphopeptides enrichment was performed as previously described(Jensen and Larsen, 2007). Briefly, stage-tips were prepared using Glass Microfibre Filters (Whatman) and 5 μm TiO_2_ beads (GL SCIENCES). Beads were suspended with 0.1% TFA in 80% AcN and inserted onto a gel-loading tip to create a 5 mm long column. Peptides samples were dissolved in 1 M Glycolic Acid 2% TFA in 80% AcN and loaded onto dedicated TiO_2_ columns by centrifugation 1 min at 6000 rpm. Columns were then washed four times with 1M Glycolic Acid 2% TFA in 80% AcN and four times with 0.1% TFA in 80% AcN. Bounded peptides were eluted first with 3% NH_4_OH and then with 0.1% TFA in 80% AcN. Eluates were then acidified with 5% TFA and dried under speed-vacuum.

#### Fe-NTA enrichment

Flow through from TiO_2_ beads were enriched using home-made stage-tips as described before but containing beads from High-Select Fe-NTA Phosphopeptide Enrichment Kit (Thermo Fisher Scientific). Enrichment was performed according to Manufacturer’s instructions.

#### ESI-LC-MSMS

Samples were diluted in 10 μl of loading buffer (5% CH3CN, 0.1% FA) and 6 μl were injected on column. LC-ESI-MS/MS was performed on an Orbitrap Fusion Lumos Tribrid mass spectrometer (Thermo Fisher Scientific) equipped with an Easy nLC1200 liquid chromatography system (Thermo Fisher Scientific). Peptides were trapped on an Acclaim pepmap100, C18, 3μm, 75μm × 20mm nano trap-column (Thermo Fisher Scientific) and separated on a 75 μm × 500 mm, C18 ReproSil-Pur from Dr. Maisch GmBH, 1.9 μm, 100 Å, home-made column. The analytical separation was run for 90 min using a gradient of H2O/FA 99.9%/0.1% (solvent A) and CH_3_CN/FA 80%/0.1% (solvent B). The gradient was run as follows: 0-5 min 95 % A and 5 % B, then to 65 % A and 35 % B in 60 min, and finally to 5% A and 95% B in 10 min with a stay for 15 min at this composition. Flow rate was of 250 nL/min. Data-dependant analysis (DDA) was performed with MS1 full scan at a resolution of 120’000 FWHM followed by as many subsequent MS2 scans on selected precursors as possible within 3 second maximum cycle time. MS1 was performed in the Orbitrap with an AGC target of 4 × 105, a maximum injection time of 50 ms and a scan range from 375 to 1500 m/z. MS2 was performed in the Orbitrap at a resolution of 30’000 FWHM with an AGC target at 5 × 10^4^ and a maximum injection time of 54 ms. Isolation windows was set at 1.6 m/z and 30% normalised collision energy was used for HCD

#### Database search

Raw data were processed using Proteome Discoverer (PD) 2.3 software (Thermo Fisher). Briefly, spectra were extracted and searched against the *Plasmodium berghei* ANKA database (PlasmoDB.org, release 38, 5076 entries) combined with an in-house database of common contaminants using Mascot (Matrix Science, London, UK; version 2.5.1). Trypsin was selected as the enzyme, with one potential missed cleavage. Precursor ion tolerance was set to 10 ppm and fragment ion tolerance to 0.02 Da. Carbamidomethylation of cysteine was specified as fixed modification. Deamidation of asparagine and glutamine, oxidation of methionine as well as phosphorylation of serine, threonine and tyrosine were specified as variable modifications. Peptide-spectrum matches were validated using the target decoy PSM validator node with a target FDR of 0.01 and a Delta Cn of 0.5. For label-free quantification, features and chromatographic peaks were detected using the “Minora Feature Detector” Node with the default parameters. PSM and peptides were filtered with a false discovery rate (FDR) of 1%, and then grouped to proteins with again an FDR of 1% and using only peptides with high confidence level. Both unique and razor peptides were used for quantitation and protein abundances are calculated as the average of the three most abundant distinct peptide groups. The abundances were normalised on the “Total Peptide Amount” and then “Protein abundance based” option was selected for protein ratio calculation and associated p-values were calculated with an ANOVA test (individual proteins).

## Notes

### Competing Interest Statement

The authors have declared no competing interest.

### Summary of Updates

Figure 1 revised; author affiliations updated.

